# Independent Influences of Movement Distance and Visual Distance on Fitts' Law

**DOI:** 10.1101/2023.09.20.558709

**Authors:** Naser Al-Fawakhiri, Samuel D. McDougle

## Abstract

Fitts’ Law is one among a small number of psychophysical laws. However, a fundamental variable in Fitts’ Law – the movement distance, D – confounds two quantities: the physical distance the effector has to move to reach a goal, and the visually perceived distance to that goal. While these two quantities are functionally equivalent in everyday motor behavior, decoupling them might improve our understanding of the factors that shape speed-accuracy tradeoffs. Here we leveraged the phenomenon of visuomotor gain adaptation to de-confound movement and visual distance during goal-directed reaching. We found that movement distance and visual distance can influence movement times, supporting a variant of Fitts’ Law that considers both. The weighting of movement versus visual distance was modified by restricting movement range and degrading visual feedback. These results may reflect the role of sensory context in early stages of motor planning.

**Public Significance:** You will automatically slow your movement when picking up a needle five inches away versus a handkerchief three inches away. This fact is elegantly formalized by Fitts’ Law, which mathematically relates movement duration to movement difficulty. However, one of the fundamental variables in the law – the distance of a planned movement – is ambiguous: Is it the actual distance the hand must move that biases movement duration, or is it the visually perceived distance? We decoupled these variables, finding that Fitts’ Law is shaped by both quantities, and that the influence of one versus the other may be related to the relevance of visual information. We believe our “addendum” to Fitts’ Law is timely, as everyday motor behavior has become increasingly enmeshed with virtual environments that abstract our movements into digital realities.

## Introduction

Motor behavior is constrained by a speed-accuracy trade-off: whenever an action must be executed quickly, its accuracy diminishes. Such a trade-off has been observed in saccadic eye movements (Gopal et al., 2017; Wu et al., 2010), goal-directed reaches (Goldberg et al., 2015), pointing movements (van Donkelaar, 1999), and even video game performance (Listman et al., 2021; Warburton et al., 2023). Fitts (1954) famously formalized the relationship between movement speed and accuracy constraints. This relationship, known as Fitts’ Law, is one of the few so-called “laws” of psychophysics and has been replicated (and occasionally caveated) extensively over decades of behavioral research in humans and other animals (Adam, 1992; de Grosbois et al., 2015; Goldberg et al., 2015; Gopal et al., 2017; MacKenzie, 1992; MacKenzie & Buxton, 1992; Sambrooks & Wilkinson, 2013; Van Gisbergen et al., 1981; Wu et al., 2010).

The common form of Fitts’ Law states that movement duration (MT) is logarithmically related to the width of a movement target (W) and the movement’s amplitude (D):

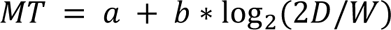

where *a* and *b* are empirically-determined free parameters. The quantity log_2_(2*D*/*W*) is typically called the Index of Difficulty (ID), reflecting a task’s accuracy constraints (other accuracy constraints, such as the weight of a held tool, can be incorporated into this index as well).

Fitts’ Law has been of particular interest in applications involving human-computer interactions (MacKenzie, 1992), such as controlling a cursor on a computer screen with a mouse (Sambrooks & Wilkinson, 2013; Thompson et al., 2004; Whisenand & Emurian, 1996), making movements in virtual or augmented reality (Rohs et al., 2011; Rohs & Oulasvirta, 2008), or performing actions in a video game (Listman et al., 2021; Warburton et al., 2023). In these virtual arenas, an inherent ambiguity in the movement amplitude (D) term of Fitts’ Law is laid bare: Does D reflect the physical distance traversed by the limb, or the perceptual distance over which one intends to move a proxy of their action (e.g., a cursor or avatar)? For example, a hand-held computer mouse only needs to move a few centimeters for its visual proxy to traverse a large computer monitor. It has been observed that as visual distance on a screen becomes increasingly decoupled from physical movement distance, predictions using Fitts’ Law tend to be worsened (Thompson et al., 2004).

Explanations for Fitts’ Law suggest that it is an outgrowth of basic facets of the motor system. For instance, it has been argued that Fitts’ Law emerges from the size of the command signal used to produce a movement – a smaller command signal may achieve greater accuracy but result in a slower movement, while a larger command signal may generate a faster but less accurate (more variable) movement. This “signal-dependent noise” model (Harris & Wolpert, 1998) suggests that movement times (MTs) reflect how the size of the motor command signal is affected by accuracy constraints, achieving a compromise between these constraints and the duration of the movement.

It has also been argued that Fitts’ Law emerges in the absence of such noisy motor commands. Al Borno and colleagues (2020) position Fitts’ Law at a more abstract level of planning: when high accuracy is demanded, many movement plans that land within the target are possible, but with a wide range of movement durations. Thus, inefficient solutions with long durations are likely to be chosen. When the accuracy constraints are relaxed, movement plans with short durations can be readily selected, but with more variable endpoints.

Beyond more fundamental aspects of action selection and the generation of motor commands, other contextual variables may influence Fitts’ Law. For instance, some have observed violations of Fitts’ Law when visual illusions alter the perception of movement amplitude (D) and/ or the width of a goal target (W), suggesting that higher-level visual representations can shape speed-accuracy tradeoffs during movement planning (van Donkelaar, 1999, but see also Alphonsa et al., 2016). These results suggest that a range of factors may contribute to how the motor system computes movement accuracy constraints and determines movement durations.

With these previous findings in mind, we hypothesized that the D term in Fitts’ Law is influenced by both visual and physical movement distances. To test this, we leveraged the phenomenon of visuomotor gain adaptation (Bock & Burghoff, 1997; Krakauer et al., 2004) to decouple movement and visual distances. We found support for our predictions in a typical reaching task: MTs in a post-adaptation, no-feedback test phase were influenced by both movement and visual distance (Exp. 1). Moreover, the relative weighting of these quantities was flexible: Visual distance dominated when the range of physical movements was restricted (Exp. 2). In contrast, when continuous visual feedback was removed during the adaptation phase, movement distance alone could explain test phase MTs (Exp. 3). These latter effects persisted when we accounted for difficulty differences between conditions attributable to latent task geometry (Exp. 4). Taken together, our results suggest that the speed-accuracy tradeoff formalized by Fitts’ Law may reflect the integration of multiple contextual variables. We speculate that this contextual influence may operate during early planning stages of goal-directed movement.

## Methods

### Participants

Experiments 1 & 4a were conducted in-lab. A total of N = 20 subjects (19 right-handed, age: 28.9 ± 9.1, 55% reported their sex as female) participated in these experiments at an honorarium of $10/hr (Exp. 1: N=10, age: 29.8 ± 10.9, 50% female; Exp. 4a: N=10, age: 27.9 ± 7.5, 60% reported their sex as female). (Sample sizes for Experiments 1 and 4a were not determined using *a priori* power analysis, though are consistent with similar investigations on the basic psychophysics of Fitts’ Law; Alphonsa et al., 2016; Rohs et al., 2011; Rohs & Oulasvirta, 2008; van Donkelaar, 1999; Wu et al., 2010). All participants provided informed, written consent in accordance with procedures approved by the Yale University Institutional Review Board, and reported their handedness using the Edinburgh Handedness Inventory, with a score of >40 indicating right-handedness (Oldfield, 1971).

Experiments 2, 3, and 4b were crowd-sourced, where data from N = 84 subjects (77 right-handed, 7 ambidextrous, age: 28.3 ± 4.6, 54% reported their sex as female) were collected online via Prolific (Exp. 2: N=27, age: 27.1 ± 5.0, 56% female; Exp. 3: N=27, age: 29.2 ± 4.7, 52% reported their sex as female; Exp. 4b: N=30, age: 28.5 ± 4.1, 53% reported their sex as female). Recruitment was restricted to right-handed or ambidextrous individuals in the United States between the ages of 18 and 35, who had at least 40 prior Prolific submissions, and a sample size of 30 was targeted for each experiment. While not based on *a priori* power analysis, this sample size mirrors typical psychophysics sample sizes.

### Apparatus

In-lab participants sat on a height-adjustable chair facing a 24.5in LCD monitor (Asus VG259QM; display size: 543.74mm x 302.62mm; resolution: 1920 x 1080 pixels; frame rate set to 240Hz; 1ms response time), positioned horizontally ∼30cm in front of the participant above the table platform, thus preventing vision of the hand **(Figure 1a)**. In their dominant hand subjects held a stylus embedded within a custom-modified paddle which they could slide across a digitizing tablet (Wacom PTH860; active area: 311mm x 216mm). Hand position was recorded from the tip of the stylus and sampled by the tablet at 200Hz. Stimulus presentation and movement recording were controlled by a custom-built Octave script (GNU Octave v5.2.0; Psychtoolbox-3 v3.0.18; Ubuntu 20.04.4 LTS).

**Figure 1.**
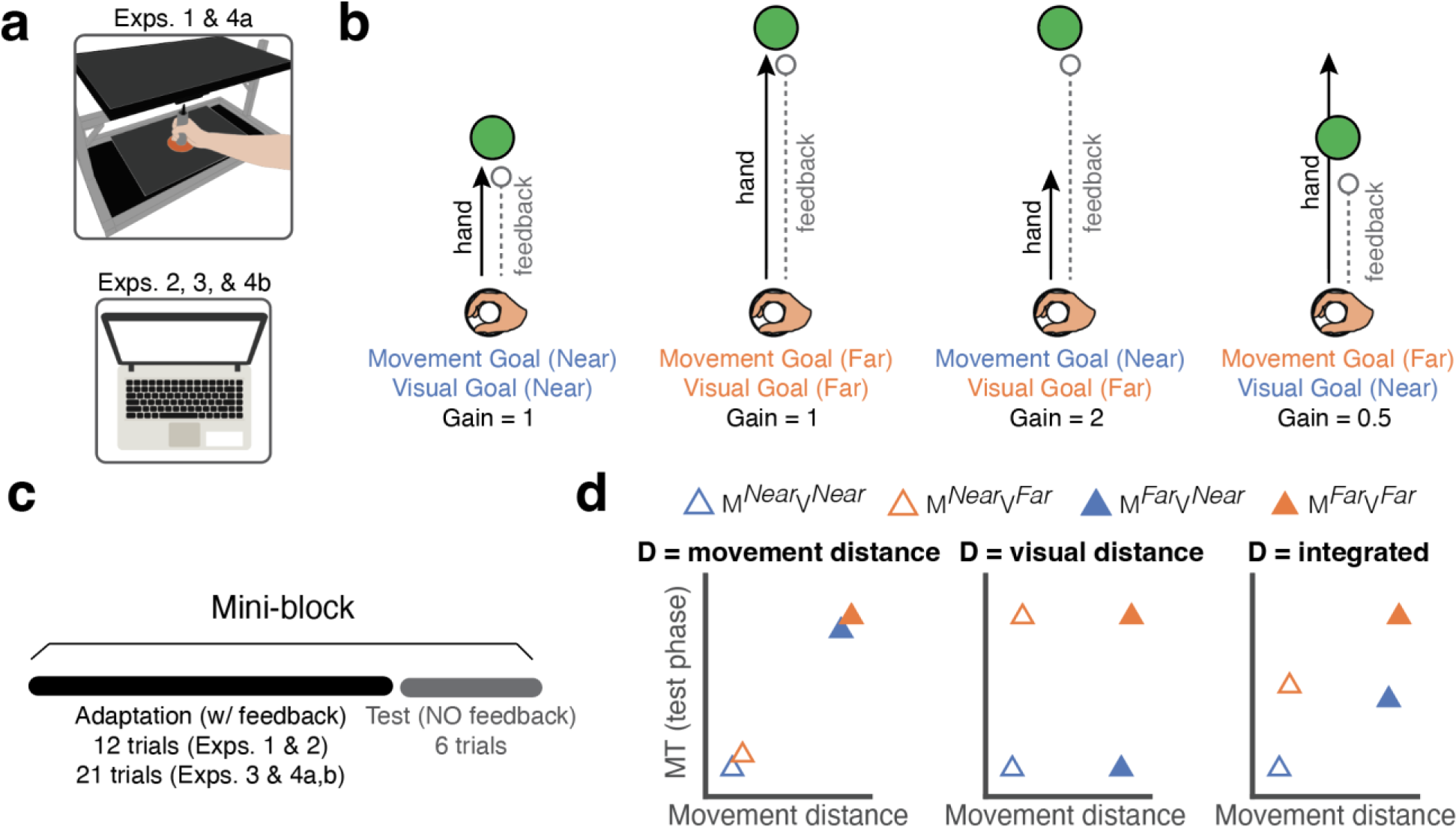
Task design and theoretical predictions. **a)** *Top:* In-lab experimental apparatus in Exps. 1 and 4A. Participants made center-out reaching movements on a digitizing tablet and viewed visual feedback on a monitor that occluded the arm. *Bottom*: Exps. 2, 3, and 4B were conducted online. **b)** Illustration of the four conditions used across all experiments – a 2x2 design crossing two levels of movement goal distances and visual goal distances by using feedback gain perturbations. **c)** Participants completed multiple mini-blocks for each of the four conditions. These mini-blocks consisted of a short adaptation phase with feedback, followed by a no-feedback test phase. Analyses were conducted on movement time (MT) in the test phase. **d)** Idealized predicted movement times given different (log) movement distances across the four conditions, for three different models. The models differ in their definition of the D term of Fitts’ Law. *Left:* movement distance (MD model); *Center*: visual distance (VD model); *Right:* multi-sensory integration of both movement and visual distance (MSI model).

### General Task Protocol

Across all experiments, participants completed center-out reaching movements and were instructed to move their hand (in-lab, **Figure 1a**) or computer mouse (online) to land a displayed cursor within a visually indicated target. Participants were instructed to always move as quickly and accurately as possible. A trial would start when participants brought their cursor to the central starting location (in-lab: 7mm diameter starting circle; online: 6 mm diameter assuming 96 PPI). To assist with re-centering, the cursor (in-lab: 3mm diameter cursor; online: 4mm diameter) was visually displayed when it was within 1cm of the start location. After waiting in the center for 500ms, a circular target would appear at one of three possible angular locations (straight ahead, or ±45° from straight ahead) and two possible distances (in-lab: 5.3cm or 10.6cm away from the start location; online: 4cm or 8cm away). For online studies, mouse acceleration was inhibited by our task code. Centimeter or millimeter distances for online studies were estimated assuming the resolution of 96 PPI (participant resolutions may have differed from this value). We note that in our online studies, we could not directly control vision of the hand during movement as we could in-lab.

We leveraged gain perturbations (Bock & Burghoff, 1997; Krakauer et al., 2004) – alterations in the relationship between movement distance and the visual consequences of those movements – to decouple physical and visual distance in “mini-blocks” of trials. We used a 2 X 2 design that yielded 4 possible conditions (**Figure 1b**): 1) the target was displayed at the shorter distance, and the required movement was similarly short (gain = 1; no perturbation), 2) the target was displayed at the farther distance, and the required movement was similarly long (gain = 1), 3) the target was displayed at the farther distance, but the required movement displacement was the same as the shorter distance (gain = 2), and 4) the target was displayed at the shorter distance but the required movement displacement was the same as the longer distance (gain = 0.5). In-lab participants would have to physically move either 5.3cm or 10.6cm for all respective trials, whereas online participants executed dramatically shorter movement amplitudes due to their use of a computer mouse or trackpad. Trials ended when participants stopped moving (velocity criterion: <5mm/s for greater than 200ms) or after a maximum time elapsed from the start of their reach. In Experiments 1 and 2, this maximum time was 1s. In Experiments 3 and 4, this maximum time was 2s.

In each mini-block (**Figure 1c**), participants first completed an “adaptation phase” (Exps. 1 & 2: 12 trials; Exps. 3 & 4: 21 trials), in which visual feedback was provided. If the visual cursor successfully landed within the target region, participants received 10 points, which was displayed adjacent to the target in green text and added to a point total displayed on the screen. Endpoint cursor feedback and points were displayed for 500ms. This adaptation phase allowed participants to learn the appropriate distances required to hit the target for each condition (i.e., to undergo gain adaptation). After adaptation, participants completed a “test phase” (all Exps.: 6 trials), in which, critically, no visual feedback nor points were provided. All key movement time (MT) analyses were performed on these testing phases to preclude effects of feedback corrections. During the testing phases, participants were instructed to keep doing what they had been doing at the conclusion of the adaptation phase, thus recreating the movements that were successful in the previous trials. Participants were informed that they would not receive any feedback about their performance in the test phase.

### Experiment 1

Participants (N=10) were provided continuous (“online”) cursor feedback during the adaptation phase of each mini-block. For trials where the visual distance and the movement distance were identical (gain = 1), the cursor veridically tracked the position of the hand. For trials where the visual distance and the movement distance were de-confounded (**Figure 1b**), the cursor was perturbed (i.e., for the condition in which the visual distance was twice as long as the movement distance, a hand displacement of 1 cm resulted in a cursor displacement of 2 cm). This manipulation is typically referred to as a gain perturbation (Bock & Burghoff, 1997; Krakauer et al., 2004), and the ratio between the visual distance and the movement distance is referred to as the gain (i.e., the previous example has gain = 2). Training mini-blocks were 12 trials in length, followed by a 6-trial test phase. Participants completed 24 training-test mini-block pairs, with 6 pairs for each condition. Conditions were pseudo-randomized such that all 4 conditions appeared in a shuffled order before repeating in a new shuffled order. The target diameter was 1.4cm and reach distances were 5.3cm or 10.6 cm.

### Experiment 2

Participants (N=27) completed a task that was largely identical to Exp. 1. The task was slightly shorter in length (16 training-test mini-block pairs; 4 per condition) to accommodate the online, crowd-sourced format. The target diameter was 60px and visual distances were 150px or 300px (assuming a standard of 96 pixels per inch, 4cm or 8cm respectively). Crucially, given the online format, the task now required only small, restricted mouse movements.

### Experiment 3

Participants (N=27) completed a similar task to Exp. 2, with some modifications. Instead of providing online feedback during the adaptation phase, feedback was degraded such that it was only provided when participants stopped moving (i.e., endpoint-only feedback). Since adaptation to endpoint feedback is slower (Taylor et al., 2014), adaptation phases were lengthened to 21 trials. Participants completed 16 training-test mini-block pairs (4 per condition). The endpoint cursor seen during adaptation followed the 2 X 2 gain perturbation design employed in Exps. 1 and 2 (**Figure 1b**). Additionally, to facilitate learning in this more difficult context, the targets were slightly larger (75 px diameter) and color-coded: blue targets indicated no gain manipulation, green targets indicated that participants needed to “go farther” than expected in order for the cursor to hit the target, and red targets indicated that participants needed to “stop shorter” than expected in order to hit the target. This instruction was repeated throughout the task at the start of each training mini-block and there was an additional 3-trial tutorial used to explain the nature of the perturbations. The target diameter was 75px and visual distances were 150 px or 300px.

### Experiments 4a and 4b

Experiments 4a (in-lab, N=10) and 4b (online, N=30) were largely identical to Exp. 3, including the 3-trial tutorial, color-coded targets, and endpoint-only feedback. Participants completed 16 training-test mini-block pairs (4 per condition) and adaptation phases were again 21 trials long. The target diameter was 2cm (75px) and reach distances were 4cm or 8cm (150px or 300px). However, in Exp. 4, the cursor was not manipulated using a “gain” manipulation. Instead, Exp. 4 aimed to preserve the “effective width” of the target across all 4 conditions. By effective width, we refer to the fact that gain manipulations will alter the area that constitutes successful reaches which would cause the cursor to land in the target. In other words, if the gain is 2 and a far target distance is displayed, any initial small deviation from the correct amplitude or angle of the reach, relative to the goal, will be magnified two-fold by the visually displayed cursor. Thus, in effect, the region which allows successful reaches to land within the targets has effectively half the radius as the equivalent (in movement distance) no-gain condition.

To control for this “geometric” difficulty difference, Exp. 4 employed a slightly modified perturbation: In conditions where the visual and movement distance were matched, no change occurred, as in the earlier experiments. However, in conditions where the movement distance was decoupled from visual distance, endpoint feedback was calculated using a novel translation that accounted for effective width (a “difficulty clamp”). In these conditions, one can imagine a virtual (invisible) target at the movement distance (where the target would have appeared on an unmanipulated trial). Reach errors were computed relative to this virtual target but were displayed relative to the visual target distance. This design maintained the basic procedures of Experiment 3 but ensured that effective target width was identical within each movement distance condition.

### Statistical Analysis

The primary dependent measure was movement duration (MT). Linear mixed-effect models were run using R’s *lmerTest* package. These models were designed to fit the MTs recorded during the test phases, using fixed effects of the physical movement extent (MD), the visual distance of the target (VD), and the task success rate (SR; i.e., the percentage of trials where subjects did not time out) during the second half of each adaptation mini-block, with subject ID as a random effect. Success rate was included as a key predictor to control for the potentially confounding effects of task success on movement speed or vigor (Summerside et al., 2018). We restricted our SR analysis to the second half of the adaptation mini-blocks to exclude errors due to relearning the perturbation and because these timepoints were closer to the critical test phases (though computing SR using the full training block did not change the results). Type III ANOVA analysis with Satterthwaite’s method was done using R’s *anova* command on the results of the LMER regression, with automatic sphericity corrections applied. It should be noted that our key analyses were only on MTs during the no-feedback test phases. Thus, these MTs were unlikely to be contaminated by corrective movements (this was further verified by examination of the trajectories, see below).

We additionally computed the physical error between the movement executed and the ideal movement. Errors were defined as the magnitude of the vector between the movement endpoint and the endpoint that would have resulted in the cursor landing in the center of the target (the optimal solution), and included trials where subjects timed out, using their last recorded hand location. Learning curves were computed using this metric, averaging across trials within each adaptation phase. We additionally defined “extent error” as the difference between the final magnitude of the reach and the optimal movement distance. Extent error was positive if participants overshot the ideal movement amplitude (regardless of the angle of the trajectory) and negative if they undershot. Success rate during the adaptation phase was also used to examine learning and was indexed by the proportion of trials in which participants landed within the target in the allotted maximum movement time for a given cycle of adaptation trials.

To target feedforward motor planning effects, post-hoc analyses of time-to-max-velocity (TTMV) were conducted on test phase reaches in Experiments 1 and 4A, the in-lab experiments. TTMV was the time from the onset of the reach until the maximum reach velocity was achieved before stopping or prior to the maximum allotted movement time. As was done with MTs, TTMVs in the test phases were fit with a linear mixed-effects regression model, where visual distance, movement distance, and success rate during the second half of each adaptation mini-block were again used as predictors. A Type III ANOVA with Satterwaithe’s method was conducted to extract the fixed effects. Additionally, we re-conducted the LMER analysis on MTs after excluding trials where the reaching angle at midpoint differed from the reaching angle at endpoint by more than 10 degrees to account for effects of corrective sub-movements. Midpoint reaching angle was defined as when the movement amplitude exceeded half of the final movement amplitude. All analyses were conducted in R (version 4.2.1).

### Modeling

Individual participant MT data were fit using various models all employing the basic Fitts’ Law formulation, MT = *a* + *b*log*(2D/W). The predicted patterns of data generated by these models are plotted in **Figure 1d**. The movement-distance-only (MD) model predicts MTs solely using the extent of the participants’ hand displacement to set the D term. The visual-distance-only (VD) model predicts MTs solely using the visually displayed distance of the target to set the D term. The integrated (INT) model employs a weighted average between the movement distance and the visual distance to set the D term on each trial:

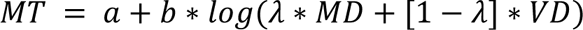

where the weighting parameter *λ* constitutes a third free parameter in addition to the standard *a* and *b* offset and scaling parameters from Fitts’ Law; this additional weighting parameter varied from 0 (only visual distance matters) to 1 (only movement distance matters).

All models assumed the target width was constant, as its visual width was identical across all conditions and experiments. As a control, we also included a fourth model that accounts for differences in “effective target widths” that are caused by gain manipulations (see descriptions for Experiment 4a and 4b above for details). This effective width (EW) model performed a weighted average between the visual width (a constant) and the effective width (the inverse of the gain applied on a particular trial):

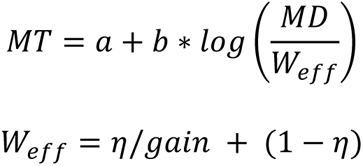

where, if the free parameter *η* is 1, the width term of Fitts’ law is entirely about the effective width of the target, and if *η* is 0, the width term is completely unaffected by the gain manipulation, reflecting the constant width of the visual target across all conditions.

Model fitting was performed using R’s *optim* function, employing the L-BFGS-B optimization method. Model comparison was conducted with the Bayesian information criterion (BIC) computed on the average of model-predicted MTs for each condition and subject. Reported “summed ΔBICs” were computed relative to the mean BIC for each subject across the three main models (INT, MD, VD). When comparing any two models, mean ΔBICs represent across-subject averages of the difference in BICs for the two models compared. Pseudo-R^2^ was computed per subject as 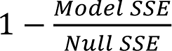 where the Model SSE was the sum of squared errors between the model-predicted average MT for each condition and the true average MT for each condition, and the Null SSE was the sum of squared errors between the true average MT for each condition and the average MT for all conditions.

### Transparency and Openness

Data and analysis scripts are available at https://zenodo.org/records/10642652. The experiments reported here were not preregistered. Ideas and data presented here were not previously disseminated.

## Results

### Experiment 1

In this study, we aimed to clarify which distance cues determine movement duration in a reaching task. Online feedback was provided and the gain on the cursor was manipulated in order to dissociate the visual distance to the target and the physical movement distance required to hit the target (**Figure 1b**). During the 12-trial adaptation phases, participants successfully adapted to the perturbation, achieving small reach errors by the last trial within each adaptation phase: The Euclidean distance from the optimal solution (i.e., the movement needed in order for the cursor to land in the center of the target) at the end of gain adaptation averaged 4.3 mm (95% CI [3.3, 5.3]). Euclidean distances (“errors”) from the optimal solution on the last trial were not significantly different between the un-manipulated (gain = 1) and manipulated (gain = 2 or 0.5) conditions (paired t-test, t(9)=-0.41, p=0.67, d_z_=-0.18). For the manipulated trials, participants slightly undershot M^far^V^near^ (signed extent error: -5.4 mm, 95% CI [-21.7, 10.9]) and significantly overshot M^near^V^far^ (22.2 mm, 95% CI [11.7, 32.8]). On average, participants successfully landed within the target on 91% of training trials (95% CI [88%, 94%]; **Figure 2b**). Success rates significantly differed between conditions (paired t-test, t(9)=3.59, p=0.005, d_z_=0.72), with manipulated trials resulting in 5.3% fewer successes on average (88.4% vs 93.7%). While statistically significant, this difference amounts to an average of one additional failed trial per 20 trials. Success rates in the second half of each adaptation block (i.e. after learning asymptote) averaged 92.0% (95% CI [89%, 95%]).

**Figure 2.**
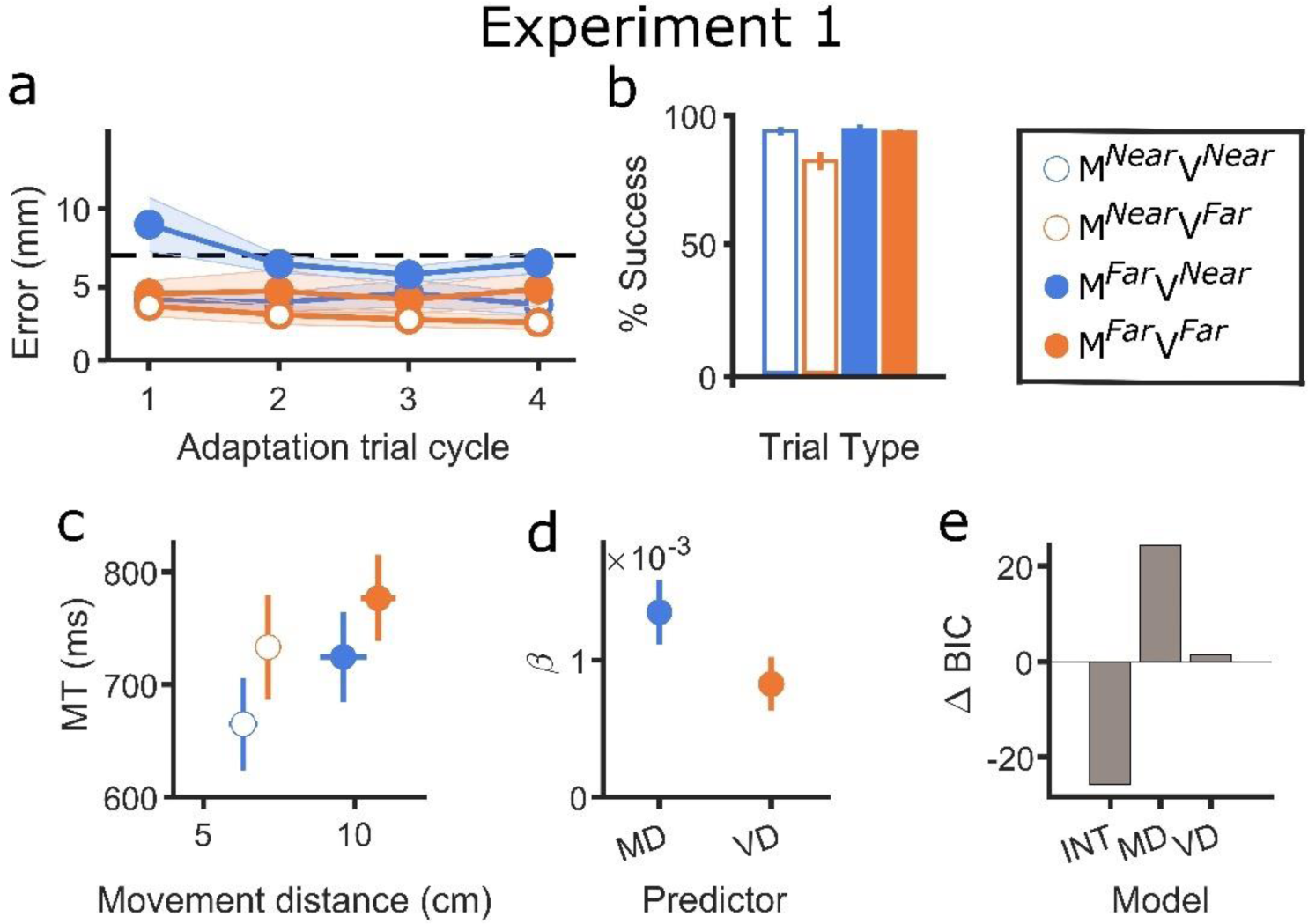
Experiment 1 results. **a)** Errors from the optimal solution (the target’s center) for each trial type during adaptation, averaged across blocks and binned per cycle. Dashed line indicates the radius of the target (7 mm). Adaptation trial cycle reflects the average of 3 reaches, one to each target location. Shaded error bars reflect standard error of the mean (SEM). **b)** Success rates during adaptation for each trial type. The middle two bars are manipulated trials. Bars on the ends are unmanipulated trials. Error bars reflect SEM. **c)** Test phase MTs plotted against the distance moved by the participant, averaged for each of the four conditions. Error bars reflect SEM for both the movement distance data (horizontal error bars) and MT data (vertical error bars). **d)** Regression coefficients for movement distance (MD) and visual distance (VD) from the LMER analysis of MTs. Error bars reflect SEM. **e)** Summed ΔBIC for the three main models (INT: Integration; MD: Movement Distance; VD: Visual Distance), computed relative to the average BIC for all three models to visualize differences. Lower values translate to stronger model fits.

Movement times during the critical test phase were fit using a linear mixed-effect model. MTs were predicted using fixed effects of true hand displacement, the visual distance of the target, the success rate for that trial type during the second half of each adaptation mini-block, and a random effect of subject ID. Actual hand displacement was used instead of ideal displacement to better capture the true data (**Figure 2c**). Success rate during late adaptation (see *Methods*) was included as a predictor to control for the effect of task success on movement vigor (Summerside et al., 2018). Crucially, an ANOVA on the regression results indicated main effects of both hand displacement (F(1, 27.39)=32.35, p=4.6x10^-6^) and the visual distance of the target (F(1, 27.14)=17.60, p=2.6x10^-4^) on test phase MTs, suggesting that both factors may influence Fitts’ Law (**Figure 2d**). However, success rate did not significantly predict test phase MTs (F(1, 27.40)=0.25, p=0.62).

These results were echoed by our computational modeling (**Figure 2e**): The INT model (summed ΔBIC=-26) outperformed both the MD (summed ΔBIC =24, mean ΔBIC=-5.00 [relative to INT], 7/10 better fit by INT) and VD models (summed ΔBIC=1.5, mean ΔBIC=-2.72, 6/10 better fit by INT). The INT model fit participant trial-type average MTs with a strong pseudo-R^2^ of 0.83 (MD: 0.40, VD: 0.54), again supporting the idea that physical distance and visual distance both contributed to MTs.

As an additional control, participant MTs were also fit with the EW model, which controls for the gain perturbation’s tendency to modify error tolerances (see *Methods*). Critically, this model failed to beat the INT model (mean ΔBIC=4.65, 8/10 better fit by INT).

Lastly, the key free parameter in the INT model (*λ*), which lineary averages the visual and movement distances, had a mean value of 0.53 (95% CI [0.28, 0.78]), indicating that roughly 53% of the D term in Fitts’ Law was reflective of the movement distance and 47% was reflective of the perceived visual distance of the target. Having demonstrated effects of both movement and visual distance on MT, we next asked how aspects of task context could alter the relative weighting of these two variables.

### Experiment 2

Experiment 2 was identical to Experiment 1 but was conducted online, allowing us to expand our sample size. The online format was chosen so that actions would be restricted to small mouse movements, significantly decreasing the range of physical movements required by each condition but maintaining large visual differences, as is typical for mouse movements in most computerized settings. It was hypothesized that this restriction might reduce the effect of the hand displacement on the index of difficulty (ID) and emphasize the effect of the visual distance to the target.

During the 12-trial adaptation phases, participants adapted to gain perturbations when they were present: By the last trial of the adaptation phases, Euclidean error from the optimal movement end-point averaged 15 pixels (25% of the target diameter; 95% CI [13, 16] px). Since Exp. 2 was conducted online, these errors reflect the difference between the participant’s unmanipulated mouse displacement and the unmanipulated mouse displacement that would have been necessary to hit the center of the target. (These errors cannot be cleanly translated into physical distances since computer mice and trackpads do not report their sensitivities for security reasons, but it should be noted that 15px is roughly 4mm on a screen using a common 96 PPI resolution.) For manipulated trials, participants slightly undershot M^far^V^near^ (extent error: -7.3 px, 95% CI [-21.7, 10.9]) and significantly overshot M^near^V^far^ (54.8 px, 95% CI [27.7, 81.9]) as in Experiment 1. Unlike in Experiment 1, Euclidean errors differed subtly between manipulated and un-manipulated trials: Errors from optimal solution on the last training trial within a block were significantly larger when the cursor gain was manipulated (paired t-test, t(26)=-4.3, p=2.4x10^-4^, d_z_=-0.92) by an average of 4.4px (95% CI [2.3, 6.6] px) (**Figure 3a**). This also corresponded to differences in success rates (**Figure 3b**): Across all training trials, manipulated trials had significantly lower success rates (paired t-test, t(26)=2.69, p=0.01, d_z_=0.54) by an average of 2.1% (95% CI [0.5%, 3.6%]). Nonetheless, participants averaged 96.8% (95% CI [95.5%, 98.0%]) success across all adaptation trials, indicating a high degree of success and rather small accuracy differences between conditions. Success rates in the second half of each adaptation block averaged 96.9% (95% CI [95.9%, 97.8%]).

**Figure 3.**
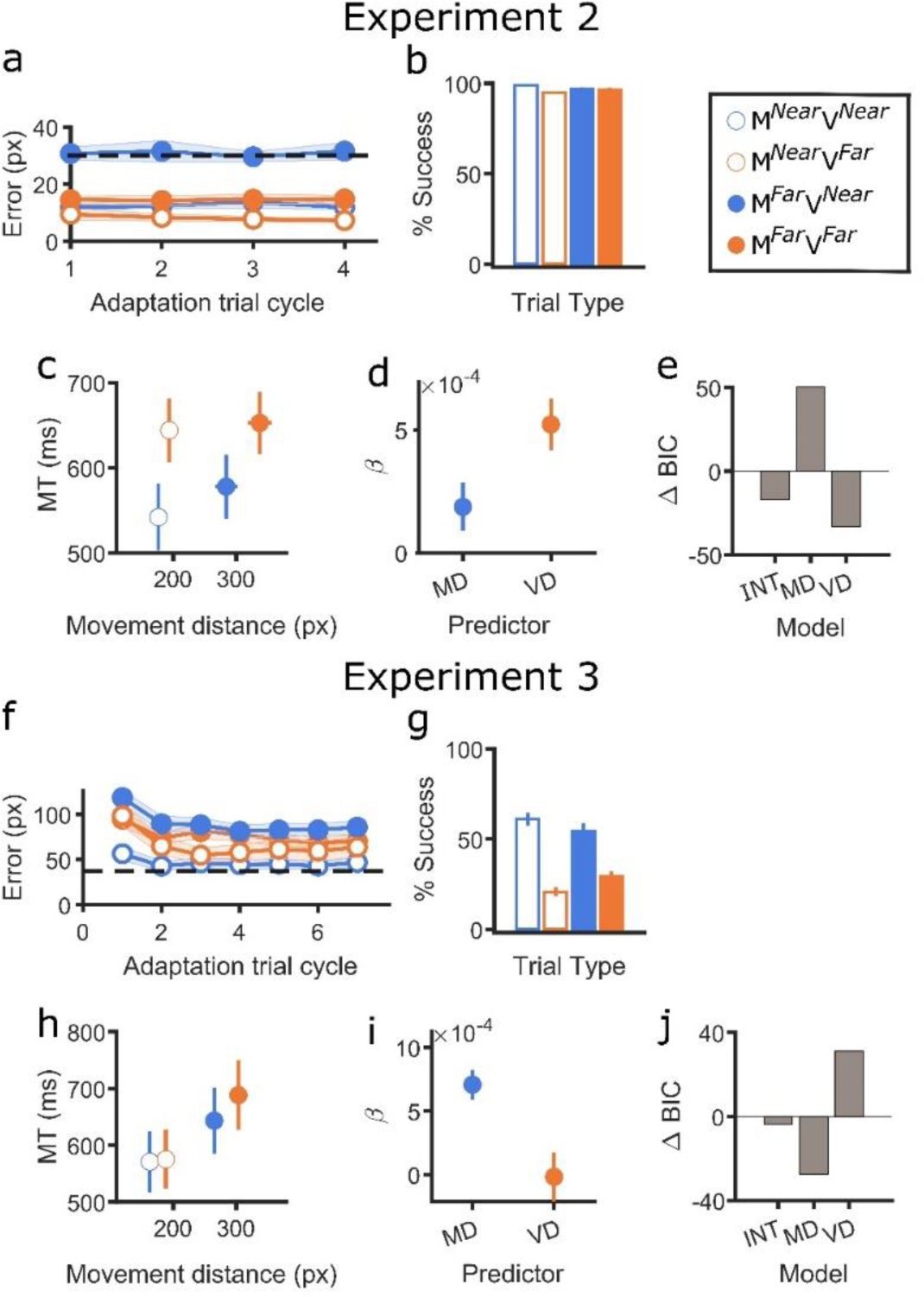
Experiment 2 and Experiment 3 results. **a, f)** Learning curves of errors from the optimal solution during the adaptation block for each condition averaged across blocks for Exps. 2 and 3, respectively. Shaded error bars reflect standard error of the mean (SEM). Dashed line indicates the radius of the target (30 and 37.5 px, respectively) **b, g)** Success rates for each trial type during adaptation. Error bars reflect SEM. **c, h)** Test phase MTs plotted against the distance moved by the participant, averaged across all four conditions for Exps. 2 and 3, respectively. Error bars reflect SEM for both the movement distance data (horizontal error bars) and MT data (vertical error bars). **d, i)** Regression coefficients for movement distance (MD) and visual distance (VD) from the LMER regression on MTs. Error bars reflect SEM **e, j)** Summed ΔBIC for the three main models (INT, MD, VD) for Exps. 2 and 3, respectively. Lower values translate to stronger model fits.

Test phase MTs were once again predicted by both movement distance and visual distance (**Figure 3c**), but not success rate during training: An ANOVA analysis over the linear mixed-effect regression on MTs showed a marginal main effect of movement (F(1, 85.20)=3.64, p=0.060) and a significant main effect of visual distance (F(1, 78.80)=25.06, p=3.3x10^-6^) (**Figure 3d**). However, success rate did not significantly predict MT (F(1, 82.99)=0.33, p=0.57). As hypothesized, due to the reduced range of the physical displacements, the effect of movement distance was reduced.

This result was further supported by modeling (**Figure 3e**): while the INT model (summed ΔBIC=-17) beat the MD model (summed ΔBIC=50, mean ΔBIC=-2.51, 15/27 better fit by INT), it did not beat the VD model (summed ΔBIC=-33, mean ΔBIC=0.59, 9/27 better fit by INT). We note that the VD model performs well here because it has fewer parameters than the INT model, and the data indicates dominance of visual distance, as hypothesized. The INT model fit participants’ trial-type MTs with a pseudo-R^2^ of 0.53 (MD: 0.19, VD: 0.46). The *λ* parameter of the MSI model, the relative weight between visual and movement distances, had a mean value of 0.50 (95% CI [0.35, 0.65]), indicating that roughly 50% of the D term in Fitts’ Law was reflective of the movement distance and 50% was reflective of the perceived visual distance of the target.

As a further control, the EW model, which controls for task geometry-induced differences in difficulty, did not win by BIC (mean ΔBIC=-0.85, 13/27 better fit by MSI, pseudo-R^2^=0.53). Having shown that we can downweight the role of movement distance and upweight the role of visual distance in Fitts’ Law, we next tried to do the opposite.

### Experiment 3

The results of Experiments 1 and 2 suggest that both movement and visual distances can influence Fitts’ Law, and that the role of visual distance can be amplified by restricting the range of possible movement distances relative to visual distances. In Experiment 3, we asked if the opposite effect could be elicited, such that movement distance could be made to primarily shape MTs, even under the restricted movement range conditions of Exp. 2. Thus, in Experiment 3 we provided only endpoint feedback during the adaptation phase. This degraded visual feedback should, we reasoned, lessen the salience of the visual aspects of the task during motor planning.

Due to the degraded visual feedback, participant errors were predictably larger in Exp. 3 compared to Exp. 2. Movement errors on the last trial of the adaptation phase were on average 67.6px (89% of the diameter of the target or roughly 1.8cm on the screen assuming 96 PPI, 95% CI [54, 81] px). Errors from the optimal solution on perturbation trials were significantly larger compared to un-manipulated trials (paired t-test, t(26)=-2.51, p=0.02, d=-0.38) by an average of 14.7px (95% CI [3, 27] px) (**Figure 3f**). For manipulated trials, participants undershot M^far^V^near^ (extent error: -23.5 px, 95% CI [-46.6, -0.3]) and overshot M^near^V^far^ (47.1 px, 95% CI [21.6, 72.5]). Manipulated trials also had lower success rates (paired t-test, t(26)=4.72, p=7.0x10^-5^, d=0.46), by an average of 7.7% (95% CI [4%, 11%]) (**Figure 3g**). Overall, success rates were substantially lower in Exp. 3 (as was expected, given the lack of online feedback), with average success across all training trials at 41.4% (95% CI [35%, 48%]) despite the lengthened adaptation phase (21 trials). Success rates in the late half of each adaptation block (i.e. after learning asymptote) averaged 43.5% (95% CI [38.6%, 48.4%]).

Participant test phase MTs were well predicted by movement distance, but, critically, not by visual distance or success rate (Figure 3h, i): ANOVA analysis revealed a main effect of movement distance on MTs (F(1, 80.09)=36.36, p=4.8x10^-8^) but no significant effect of visual distance (F(1, 85.38)=0.01, p=0.93) nor success rate (F(1, 87.73)=0.03, p=0.85).

Model comparison (**Figure 3j**) using BIC indicated that the MD model (summed ΔBIC=-28) was the best-fitting model (compared to INT [summed ΔBIC=-3]: mean ΔBIC=-0.88 [relative to MD], 24/27 better fit by MD; compared to VD [summed ΔBIC=31]: mean ΔBIC=-2.18, 20/27 better fit by MD). However, the INT model had the highest pseudo R^2^ at 0.46 (MD: 0.41, VD: 0.12). The INT model’s ability to capture more of the variance in MTs did not overcome the BIC penalty for an additional parameter. The *λ* parameter, which estimates the relative contribution of visual and movement distances on D, had a mean value of 0.76 (95% CI [0.64, 0.87]), indicating that roughly 76% of the D term in Fitts’ Law was reflective of the movement distance and 24% was reflective of the perceived visual distance of the target. The control EW model was largely equivalent to the INT model (mean ΔBIC=-0.16 in favor of EW, 13/27 better fit by EW).

These observations support the conclusion that MTs in Exp. 3 were primarily determined by movement distances as opposed to visual distances, in stark contrast to Exp. 2. Given that the underlying gain manipulation in Exp. 3 was identical to Exps. 1 and 2, and that the movement demands fully matched Exp. 2, the flip in the factor shaping MTs from visual to movement distance appeared to be driven solely by the change in the visual feedback context. Taken as a whole, the results of Exps. 1-3 point to movement and visual distance independently influencing the D term in Fitts’ Law.

### Experiment 4a

Experiments 1-3 employed conventional gain perturbations in which small displacements, Δ*x*, are displayed as Δ*x* ∗ *gain*. As a result of this, the “effective width” (that is, differences in motor error tolerance in gain versus non-gain contexts; see *Methods*) of the movement targets varies across conditions with different gains, in that gain can inflate or deflate errors in initial movement direction and extent. (For example, when gain = 2, any slight deviation from the correct movement amplitude would be magnified two-fold in the observed visual error.) To control for this geometric difference, which is inherent to gain perturbations, Experiments 4a and 4b employed a novel translational perturbation where the visual feedback was “teleported” along the straight-line path to the target to either a greater or shorter amplitude, depending on the gain (**Figure 4a**). The size of the effective area of successful reach vectors was thus standardized for all conditions.

**Figure 4.**
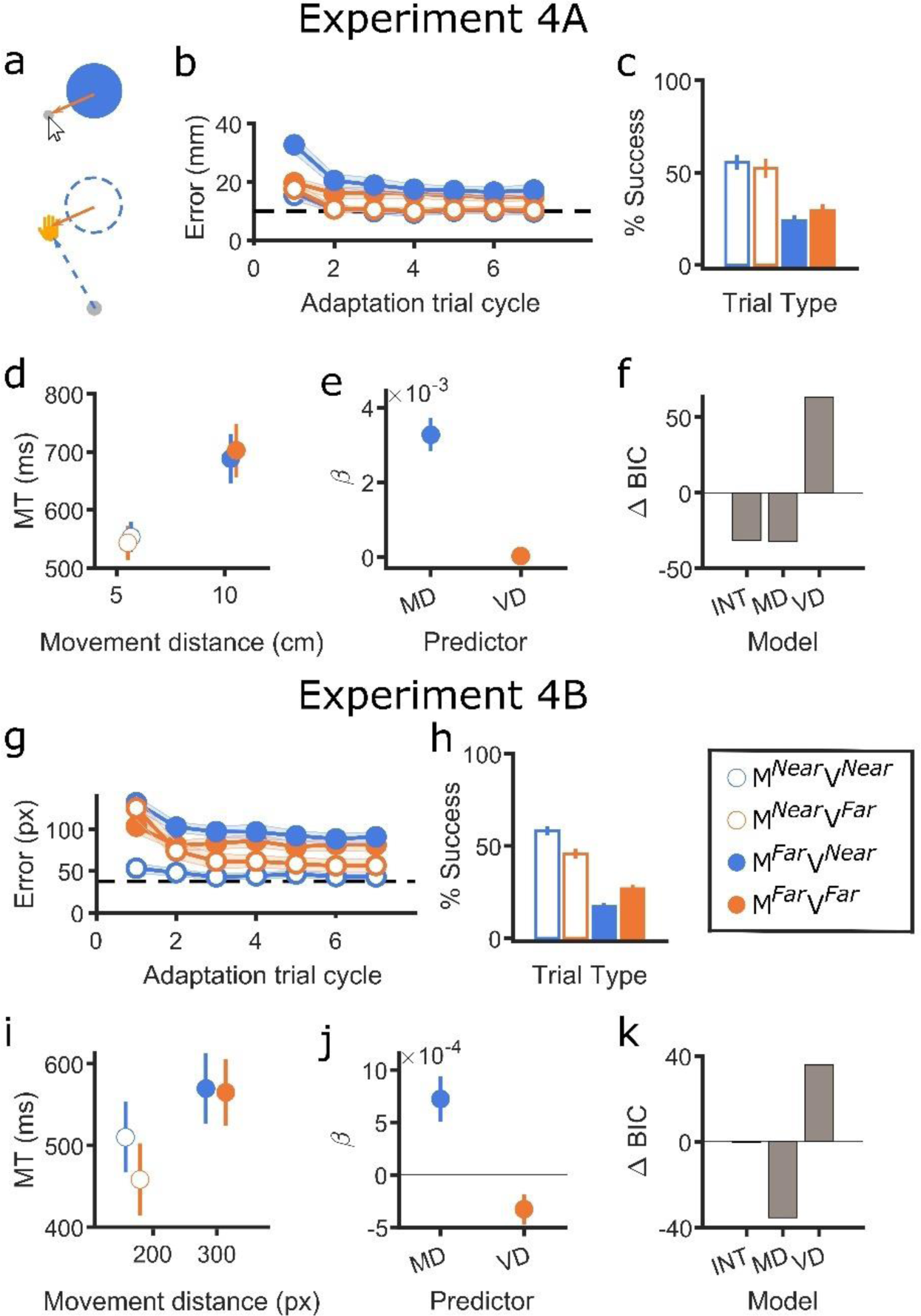
Experiment 4 results. **a)** Modified gain perturbation employed in Exp 4. Participants reaches were evaluated as though the target were at the movement distance for that trial type. Errors from that virtual target (dashed) were then displayed at the visual distance (filled). **b, g)** Learning curves of errors from the optimal solution during the adaptation block for each condition averaged across blocks for Exps. 4A and 4B, respectively. Shaded error bars reflect standard error of the mean (SEM). Dashed line indicates the radius of the target (10 mm and 37.5 px, respectively) **b, g)** Success rates for each trial type during adaptation. Error bars reflect SEM. **c, h)** Test phase MTs plotted against the distance moved by the participant, averaged across all four conditions for Exps. 4A and 4B, respectively. Error bars reflect SEM for both the movement distance data (horizontal error bars) and MT data (vertical error bars). **d, i)** Regression coefficients for movement distance (MD) and visual distance (VD) from the LMER regression on MTs. Success rate, the other predictor in the model, is not depicted since it does not share the units of distance. Error bars reflect SEM. **e, j)** Summed ΔBIC for the three main models (INT, MD, VD) for Exps. 4A and 4B, respectively. Lower values translate to stronger model fits.

Due to the discrete nature of this perturbation, only endpoint feedback could be provided. Despite this degraded feedback, participants learned to adequately adapt to the perturbation during gain trials, achieving average errors of 13.3mm (67% of target diameter; 95% CI [11, 16] mm) on the last trial of the adaptation phase. Errors from the optimal solution were not significantly different between manipulated and un-manipulated trials (paired t-test, t(9)=-0.86, p=0.41, d=-0.33). Extent errors averaged -0.2 mm (95% CI [-7.1, 6.8]) for M^far^V^near^ and 4.0 mm (95% CI [0.7, 7.3]) for M^near^V^far^, a slight overshoot. Success rates during training were also not significantly different between manipulated trials and un-manipulated trials (paired t-test, t(9)=2.16, p=0.06, d=0.39). Overall, success rates were low, averaging 40.3% (95% CI [32%, 48%]). Success rates in the late half of each adaptation block (i.e. after learning asymptote) averaged 43.4% (95% CI [36.9%, 50.0%]).

Even with a rather different perturbation applied, the test phase results of Experiment 4a largely replicated the results of Experiment 3 (**Figure 4b**). The ANOVA analysis revealed a main effect of movement distance (F(1, 29.57) = 53.60, p=4.1x10^-8^) but not visual distance (F(1, 26.92) = 0.01, p=0.92) on test phase MTs (**Figure 4c, d**). As in the previous experiments, success rate did not significantly predict MT (F(1, 30.71)=0.30, p=0.59), further suggesting that success during training did not explain differences in movement vigor during the test phase.

Modeling further corroborated these results (**Figure 4e**): By BIC comparison, the INT (summed ΔBIC=31) and MD (summed ΔBIC=32) models were largely identical in fit quality (mean ΔBIC=-0.08 in favor of MD, 6/10 better fit by MD). However, the INT model had a pseudo-R^2^ of 0.89, as opposed to the MD model at 0.79. The average value of the fitted *λ* parameter was 0.91 (95% CI, [0.83, 0.99]), indicating that movement distance contributed ∼91% of the D term in Fitts’ Law. Thus, even after accounting for the effect of varying effective target widths and altering the pattern of adaptation phase errors across conditions relative to Exp. 3, the key results of Exp. 3 were replicated in Exp. 4a.

### Experiment 4b

We next replicated Exp. 4a in an online sample. As with Exp. 3, the degraded visual feedback led to relatively large errors on the last trial of the adaptation phase (**Figure 4g**), averaging 67.7px (89% of the diameter of the target; 95% CI [58, 78] px; roughly 1.8 cm assuming 96 PPI). Errors from the optimal solution were marginally different between manipulated and un-manipulated trials (paired t-test, t(29)=-2.00, p=0.054, d=-0.33), with errors 10.5px (∼2.8mm assuming 96 PPI; 95% CI [0, 21] px) larger for manipulated trials. Extent errors averaged -0.1 px (95% CI [-26.3, 26.1]) for M^far^V^near^ and 40.1 px (95% CI [21.3, 58.9]) for M^near^V^far^, overshooting by just over one target radius. Un-manipulated trials had significantly higher success rates (paired t-test, t(29)=9.32, p=3.2x10^10^, d=0.96), by an average of 11% (95% CI [9%, 13%]) (**Figure 4h**). The average success rate across all training trials was 37.0% (95% CI [33.3%, 40.6%]). The average success rate across training trials in the second half of each adaptation block was 40.1% (95% CI [36.3%, 44.0%])

The ANOVA revealed a main effect of movement distance, as expected (F(1, 95.36)=10.93, p=1.3x10^-3^), and a significant main effect of visual distance (F(1, 88.02)=5.29, p=0.02). However, the longer visual distances actually predicted *lower* MTs (**Figure 4j**), an effect that was largely driven by faster MTs on the condition where the movement goal was near, but the visual target was far (**Figure 4i**). Interestingly, for this condition, when compared to the un-manipulated near-movement condition, participants had marginally larger errors (t(29)=1.77, p=0.09, d=0.38; 13.8px, 95% CI [-2, 30] px) and significantly lower success rates (t(29)=6.76, p=2.04x10^-7^, d=0.86; 12%, 95% CI [9%, 16%]) during adaptation phases, yet had faster MTs during the test phase (t(29)=-2.40, p=0.02, d=-0.22; 52ms, 95% CI [8, 96] ms). There was not a significant main effect of training success rate on test phase MTs (F(1, 97.69)=0.65, p=0.42).

Modeling supported the conclusion that MTs were predominantly driven by movement distance (**Figure 4k**): The MD model (summed ΔBIC=-35) won model comparison by BIC, beating the INT model (summed ΔBIC=0, mean ΔBIC=-1.17 [relative to MD], 27/30 better fit by MD) and the VD model (summed ΔBIC=36, mean ΔBIC=-2.37, 22/30 better fit by MD). The INT model had a slightly higher pseudo-R^2^ compared to the MD model (MSI: 0.38, MD: 0.36, VD: 0.05). We note that the lower pseudo-R^2^ values in general arise from the models’ inability to account for the aforementioned violation of Fitts’ Law in one of the conditions. The average value of the fitted *λ* parameter was 0.79 (95% CI, [0.68, 0.91]), indicating that movement distance contributed 79% of the D term in Fitts’ Law.

Overall, the key test phase findings of both Exps. 4A and 4B did not differ from the results of Experiment 3 – this echoes the poor fit of the EW model in the previous studies, and further suggests that the key results of our study were likely not driven by differences in target effective widths nor other difficulty considerations. Instead, the four experiments suggest that both movement distance and visual distance can influence Fitts’ Law.

#### Estimating the role of visual and movement distances on initial movement plans

Experiments 1-4 demonstrate that, under different conditions, movement distance and visual distance to a goal can have varying roles on movement durations. Specifically, when movement range was reduced, visual distance seems to exert a greater role on MTs, and, when visual feedback is degraded, movement distance exerts a greater role on MTs. How much of the observed MT effects can be ascribed to planning factors initialized before the start of movement execution?

To assess this, we performed a post-hoc analysis on time-to-max-velocity (TTMV) during the no-feedback test phases. We restricted our analysis to only the in-lab experiments (Experiments 1 and 4A), where we could obtain a reliable estimate of TTMV. As was done with the MTs, we fit a linear mixed-effects regression model on the TTMVs, with movement distance, visual distance, and the success rate in the second half of each adaptation mini-block. Again, success rate was included as a predictor to control for the effect of success on movement vigor. Fixed effects were extracted using a Type III ANOVA, as above.

In Experiment 1, only the visual distance of the target predicted TTMVs (F(1, 27.76)=4.81, p=0.04), but not movement distance (F(1, 29.23)=1.12, p=0.30) or success rate (F(1, 29.28)=0.02, p=0.90). Thus, initial movement vigor in Experiment 1, where continuous visual feedback was provided during the preceding adaptation phase, was only reliably predicted by visual target distance. The lack of a significant effect of movement distance could be because TTMV was a noisier metric than MT or that initial movement vigor only reflects planning components related to visual inputs, while movement distance exerts its effect on MT during the deceleration phase. However, in Experiment 4A, only movement distance to the target predicted TTMV (F(1, 30.58)=15.12, p=5.1x10^-4^), but neither visual distance (F(1, 26.97)=0.12, p=0.73) nor success rate (F(1, 32.09)=0.06, p=0.81).

As an additional control, we re-conducted the main LMER analyses on movement durations after excluding all trials where the reaching angle at midpoint differed from the reaching angle at endpoint by +/-10° or more. This control analysis aimed to eliminate any trials where corrective moments were suspected, though may also have excluded particularly curved ballistic reaches. This analysis excluded only a small proportion of test phase trials across all experiments (Exp. 1: 3.1%, 95% CI [1.3%, 4.9%]; Exp. 2: 8.6%, 95% CI [6.1%, 11.1%]; Exp. 3: 8.0%, 95% CI [3.7%, 12.3%]; Exp. 4A: 1.6%, 95% CI [0.3%, 12.9%]; Exp. 4B: 8.4%, 95% CI [4.0%, 12.8%]), suggesting that movements were generally rather straight. Excluding these trials changed none of the main results of the LMER analysis on MTs across all experiments. Thus, we did not observe evidence that corrective movements in the test phase played a role in explaining our results. Taken together, our analysis and velocity analyses above complement our earlier MT analyses, reflecting how visual and movement distances can independently shape movement planning processes.

## Discussion

What information does the brain use to predict the difficulty of a movement? Across four experiments we sought to clarify the nature of the movement distance (D) term of Fitts’ Law (**Figure 1d**). We leveraged gain adaptation (**Figure 1b**) to create conditions where both visual and **motor** aspects of a reaching task could independently influence movement times (MTs). Our results showed that when continuous visual feedback is provided during adaptation (Exps. 1 & 2), the visually perceived distance of a goal and the reach amplitude needed to reach that goal both contributed to MT (**Figures 2 and 3**). The role of vision was especially pronounced when movement amplitude was restricted (Exp. 2; **Figure 3**). However, when visual feedback was degraded to only appear at the endpoint of a movement, visual goal distance no longer predicted MT even within the same restricted movement context (Exp. 3; **Figure 3**). When potential condition difficulty differences due to the geometry of the task were accounted for (Exp. 4a & b; **Figure 4**), these latter results remained largely unchanged, and task success during the adaptation phase was unable to predict the key test phase MT results across all experiments. Crucially, all key results were quantified in a test phase where visual feedback was withheld; thus, differences in MTs could not be easily explained by online, corrective movements or similar reactive strategies, a conclusion that was further supported by analyses that focused on initial movement vigor or that excluded suspected corrective movements.

What is the underlying process that explains our findings? Across all experiments, success rates during adaptation were not predictive of test phase movement durations, so we do not believe that success or reward was the primary driver of the observed movement time effects. Moreover, our analyses were constrained to a test phase without visual feedback and were generally replicated when restricting our analysis to an early phase of movement: this supports the idea that the distance effects are relevant to motor planning, not feedback control. We believe that our results suggest that visual distance cues, in conjunction with past movement experiences, may provide a contextual reference during movement planning, perhaps by modulating the motor system’s expectations of movement difficulty. When movement range is limited, visual representations may come to dominate (Exp 2). When visual feedback is degraded (Exp 3), the effect of visual distance may be diminished due to a fundamental change in the planned control policy, where the visual correlates of movement plans are now less relevant and purely motor variables may play a more pronounced role.

We suggest that visual distance may exert its effect during an early motor planning stage (Al Borno et al., 2020). The visual distance to the goal, even when decoupled from the required movement distance, may act as a “prior” on the expected difficulty of a movement. Alternatively, it is also possible that larger visual distances may also increase the sensory uncertainty at the target location, as the target moves further out in the periphery relative to the starting hand location. This additional uncertainty may lead to higher difficulty expectations, and thus slow down MTs. Whether visual target distance or uncertainty (or both) drive our observed effects is unclear – future studies could attempt to isolate spatial distance effects versus spatial uncertainty effects in Fitts’ Law.

Another open question is the role of proprioceptive/kinesthetic feedback in our results. It could be that, during the adaptation phase, participants weighed kinesthetic input more heavily in Experiments 3 and 4 (when only terminal endpoint feedback was provided), perhaps explaining why test phase movement durations in these experiments were better predicted by movement, rather than visual, distance. Indeed, the distinct behavioral effects of online versus endpoint feedback we observed here echo findings in motor adaptation that show similar tradeoffs between vision and proprioception in online (visual emphasis) versus endpoint (proprioceptive emphasis) feedback conditions (Hayashi et al., 2020; Izawa & Shadmehr, 2011). Supporting a role for proprioception here, past studies on a deafferented subject showed that proprioception directly influences the relationship between movement duration and movement amplitude (Ingram et al., 2000). How could differences in the weighting of proprioception and vision influence feedforward motor planning and MTs in our task? Speculatively, this could be driven by top-down attention to each sense, which leads to a bias in MTs that is carried over to the test phase – Attention to proprioception may bias motor planning toward representations in intrinsic body-based coordinates, while attention to visual outcomes (say, attention to the online cursor) may bias planning toward representations in extrinsic world-based coordinates. These internal and external reference frames of planning may contribute to differences in MTs as a function of visual or movement distance, as observed in our experiments. Future investigations should directly measure the contributions of proprioception to the results observed here.

We think that it is also worth considering that our gain perturbation may be compensated for by a combination of explicit and implicit motor learning processes (Krakauer et al., 2004; McDougle et al., 2016; Taylor et al., 2014). That is, it is currently unclear if people compensated for the gain perturbations implicitly, explicitly, or in some combination of the two. Depending on how they learned, different interpretations of the test phase results could be favored (i.e., focusing on conscious selection processes versus implicit planning processes). Future work could target the role of explicit cognitive processes versus implicit processes in our task, and in Fitts’ Law more generally.

We highlight several limitations in our study. First, gain perturbations inherently alter error “tolerance,” as similar movement commands can result in different magnitudes of visual error. However, we think that it is unlikely that test phase MTs were significantly impacted by the latent changes in task geometry due to gain perturbations: When we directly controlled for these intrinsic condition difficulty differences in Exp. 4, the results of Exp. 3 largely replicated. While this does not fully rule out a potential role of the ‘effective width’ (see *Methods*) of reach targets in our task, in Exps. 1-3, vastly different visual and movement distance effects were observed even when effective widths did not differ across the experiments.

Second, in general, when task performance during the adaptation phase was relatively weaker, test phase MTs were slower, raising concerns of task success acting as a confound. This result makes some sense (Adam, 1992; de Grosbois et al., 2015; Fitts, 1954; Fitts & Peterson, 1964; van Donkelaar, 1999) – Fitts’ Law relates the difficulty of a movement to its duration. However, we did not always see a 1-1 relationship between task success during the adaptation phase and the key MT results in the testing phase. More importantly, all of our key regression results controlled for effects of task success. It can be difficult to fully separate effects of accuracy and movement time (their close relationship is the point of Fitts’ Law), and our study targeted MT effects following a gain adaptation phase that may involve additional difficulty variables not accounted for in our models. For example, in Experiments 1 and 2, participants may have learned to associate particular visual distances with the ease of cursor control and the cursor speed. However, if these associations persisted into the probe phase where no feedback was provided, this should have been revealed in our analyses of error and task success, but success rate during adaptation did not predict MTs when movement and visual distance were also included as model predictors. Thus, we believe that our results are parsimoniously explained by our computational models targeting the D term of Fitts’ Law. Future studies could further explore these and other aspects of difficulty.

More broadly, we think our results could contribute to a synthesis of documented amendments to Fitts’ Law (Crossman & Goodeve, 1983; de Grosbois et al., 2015; Heath et al., 2011). First, our findings are consistent with literature suggesting that strategic attention to a range of task dimensions shape Fitts’ Law (Adam, 1992). Others have also noted that past movement history can influence movement durations in a manner that is not consistent with Fitts’ Law (Tang et al., 2018). Our results also echo and extend findings that Fitts’ Law can emerge at perceptual or mental “simulation” stages of motor planning (Decety & Jeannerod, 1995; Grosjean et al., 2007), are consistent with findings that visual illusions can bias Fitts’ Law (van Donkelaar, 1999), and comport with studies relating the law to more abstract features of motor preparation and planning (Augustyn & Rosenbaum, 2005; Jax et al., 2007). Incentives also modulate movement durations (Ashworth-Beaumont & Nowicky, 2013; Bogacz et al., 2010; Du et al., 2022; Listman et al., 2021; Thura et al., 2014), further supporting flexibility in the strategies the brain employs when selecting movement parameters. We also note that other studies of Fitts’ Law in virtual environments, where visual and physical dimensions of the target do not necessarily match, largely align with our results in showing that both the visual and motor dimensions can predict movement durations under specific contexts (Usuba et al., 2019, 2021). Future investigations can address other, perhaps more cognitive indicators of difficulty, such as the visually inferred weight or unwieldiness of a tool (Ellis & Lederman, 1993).

As interest in the applicability of Fitts’ Law to virtual and augmented reality grows (Rohs et al., 2011; Rohs & Oulasvirta, 2008; Sambrooks & Wilkinson, 2013), it is important to understand the law’s underlying mechanisms. We demonstrated here that under certain conditions, fundamental variables in Fitts’ Law are responsive both to perceptual **context** (visual goal distance) and physical requirements (movement distance). Developers of AR/VR applications could leverage this observation in optimizing or calibrating the feedback they provide to users. Our results may also inform investigations into the neural correlates of action selection and the speed-accuracy trade-off (Al Borno et al., 2020; Harris & Wolpert, 1998), perhaps serving to illuminate distinct contextual factors that the motor system represents as it prepares fast and accurate actions.

## Constraints on Generality

Fitt’s Law is a surprisingly robust psychophysical law with broad applicability. Samples reported here may not be generally representative of the global population as participants were recruited from the US and were between the ages of 18-40, and were largely right-handed. Nonetheless, we believe these results should broadly apply to all neurotypical adults, as they relate to the typical performance of goal-directed movements.

